# SPRITE: improving spatial gene expression imputation with gene and cell networks

**DOI:** 10.1101/2024.01.31.578269

**Authors:** Eric D. Sun, Rong Ma, James Zou

## Abstract

Spatially resolved single-cell transcriptomics have provided unprecedented insights into gene expression *in situ*, particularly in the context of cell interactions or organization of tissues. However, current technologies for profiling spatial gene expression at single-cell resolution are generally limited to the measurement of a small number of genes. To address this limitation, several algorithms have been developed to impute or predict the expression of additional genes that were not present in the measured gene panel. Current algorithms do not leverage the rich spatial and gene relational information in spatial transcriptomics. To improve spatial gene expression predictions, we introduce SPRITE (Spatial Propagation and Reinforcement of Imputed Transcript Expression) as a meta-algorithm that processes predictions obtained from existing methods by propagating information across gene correlation networks and spatial neighborhood graphs. SPRITE improves spatial gene expression predictions across multiple spatial transcriptomics datasets. Furthermore, SPRITE predicted spatial gene expression leads to improved clustering, visualization, and classification of cells. SPRITE is available as a software package and can be used in spatial transcriptomics data analysis to improve inferences based on predicted gene expression.

## 1 Introduction

The advent of spatially resolved single-cell transcriptomics have provided new opportunities to study the biological processes governing cellular interactions and the organization of tissues *in situ* [1]. Although these spatial transcriptomics methods can detect transcripts at single-cell resolution, they are generally limited to measurement of a small number of genes [2]. Due to the resource-intensive nature of current spatial transcriptomics technologies, it is often infeasible to measure additional genes in a spatial transcriptomics experiment and as such, computational methods for predicting the spatial expression of additional genes of interest are desirable.

Several computational methods have been developed to impute or predict spatial gene expression for spatial transcriptomics datasets by leveraging whole-transcriptome gene expression information from paired single-cell RNAseq datasets. These approaches typically involve jointly embedding the spatial and RNA-seq data. After this joint embedding, these methods then predict the expression of new genes by aggregating information across neighboring cells in the RNAseq data [3, 4, 5, 6]. Prediction methods employing other methods, optimal transport for example [7], have also been proposed. In virtually all cases, important relational information such as that between genes (e.g. co-expression) and that between cells (e.g. spatial proximity) are not explicitly utilized in predicting spatial gene expression. Incorporating additional relational information in these prediction methods may provide one avenue to improve upon the current state-of-the-art in prediction of spatial gene expression. More accurate predictions of spatial gene expression are desirable and can likely improve the quality of downstream inference that rely on these predictions.

Here we develop and introduce SPRITE (Spatial Propagation and Reinforcement of Imputed Transcript Expression), a meta-algorithm that works with any existing spatial gene expression prediction method. SPRITE employs a two-step approach that centers around information propagation in gene correlation networks and spatial neighborhood graphs to refine the baseline predicted spatial gene expression obtained from an existing method. We show that post-processing of spatial gene expression predictions with SPRITE generally leads to more accurate predictions and that these improvements translate to improved performance on common downstream analysis tasks for spatial transcriptomics.

## 2 Materials and Methods

### 2.1 Benchmark datasets

We evaluated the performance of the SPRITE across eleven benchmark spatial transcriptomics and RNAseq dataset pairs [2, 8, 9, 10, 11, 12, 13, 14, 15, 16, 17, 18, 19, 20, 21, 22, 23, 24, 25] that were compiled and processed by earlier studies [2, 26], which is where they can be accessed. The spatial transcriptomics and RNAseq dataset pairs were chosen from the same species and approximately the same tissues. The dataset pairs spanned four organisms (human, mouse, fruit fly, axolotl); eight spatially resolved single-cell transcriptomics technologies, multiple single-cell RNAseq technologies, and multiple tissues. Before spatial gene expression prediction, the counts in the RNAseq datasets were normalized such that the total count for each was equal to the median of total counts across cells, and then the normalized counts were log-transformed with an added pseudocount.

### 2.2 Spatial gene expression prediction

As a meta-algorithm, SPRITE uses the baseline predicted expression obtained from an existing spatial gene expression prediction method. We evaluated SPRITE on three spatial gene expression prediction methods: SpaGE [3], Tangram [7], and Harmony-kNN [27]. SpaGE performs alignment using domain adaptation and subsequent prediction by *k*-nearest neighbors regression [3]. We used at least 20 and up to half of the principal vectors for SpaGE prediction. Tangram maps RNAseq expression onto space using a deep learning approach. We performed additional preprocessing steps for the input data according to the suggested approach in Tangram [7]. Harmony-kNN uses Harmony [27] to jointly embed the spatial transcriptomics and RNAseq data and then predicts expression for a cell in the spatial transcriptomics data by averaging gene expression from its *k* nearest neighbors in the RNAseq data. We used *k* = 10 and up to the first 30 harmonized principal components.

### 2.3 SPRITE meta-algorithm

The inputs to the SPRITE meta-algorithm are paired single-cell datasets from spatial transcriptomics and RNAseq and a spatial gene expression prediction method (see above section for examples). The goal of SPRITE is generate predictions of spatial gene expression for a set of genes that are not seen in the spatial transcriptomics data but are present in the RNAseq data. We denote the spatial transcriptomics data *X*_spatial_ ∈ ℝ^*n×p*^ and the RNAseq data *X*_rna_ ∈ ℝ^*m×q*^. In both matrices, the rows correspond to cells and the columns correspond to genes. In most cases where spatial gene expression prediction is desirable, it is true that *q* ≫ *p* (i.e. there are many times more genes in the RNAseq data as in the spatial transcriptomics data) and the genes in *X*_rna_ superset the genes in *X*_spatial_. In practice, the latter attribute can be enforced by subsetting the spatial transcriptomics data to only include genes also present in the RNAseq data.

The spatial gene expression prediction problem involves predicting the expression of a gene indexed by *j* in *X*_rna_ that is not present in *X*_spatial_. Using a spatial gene expression prediction method, the expression of gene *j* for each cell in *X*_spatial_ is predicted using information jointly obtained from *X*_spatial_ and *X*_rna_. We denote the matrix containing the predicted gene expression for *r* genes as *G* ∈ ℝ^*n×r*^.

The spatial gene expression prediction workflow including the SPRITE meta-algorithm and its steps are depicted in Fig. 1. Given a black-box gene expression prediction method (e.g. SpaGE, Tangram, or Harmony-kNN) and paired datasets *X*_spatial_ and *X*_rna_, we generate predicted expression for a set of “target” genes that are present in *X*_rna_ but not in *X*_spatial_ and also a set of “calibration” genes that are present in both *X*_rna_ and *X*_spatial_. The calibration genes are used to estimate prediction errors. To predict calibration gene expression without exposing the prediction method to the measured expression of these genes, we used a 10-fold cross-validation approach where for each fold, we excluded approximately 10% of calibration genes and predicted their expression using the remaining genes. This procedure is repeated until predictions are made for all calibration genes in the spatial transcriptomics data. After combining the predicted expression of the target genes and calibration genes into a single matrix *G*, SPRITE then performs a two-step post-processing procedure to improve upon the initial predictions. First, SPRITE reinforces the prediction errors generated using the calibration genes by propagating these errors across a gene correlation network to correct predictions for target gene expression. We refer to this first step as the “Reinforce” step. Next, the predicted gene expression is smoothed across all cells in a spatial neighborhood graph such that neighboring cells with similar measured gene expression (i.e. of the same cell type or subtype) will tend to have more similar predicted gene expression. We refer to this second step as the “Smooth” step. Finally, the SPRITE predicted spatial gene expression can be used in downstream analysis tasks that rely on gene expression data.

**Figure 1:**
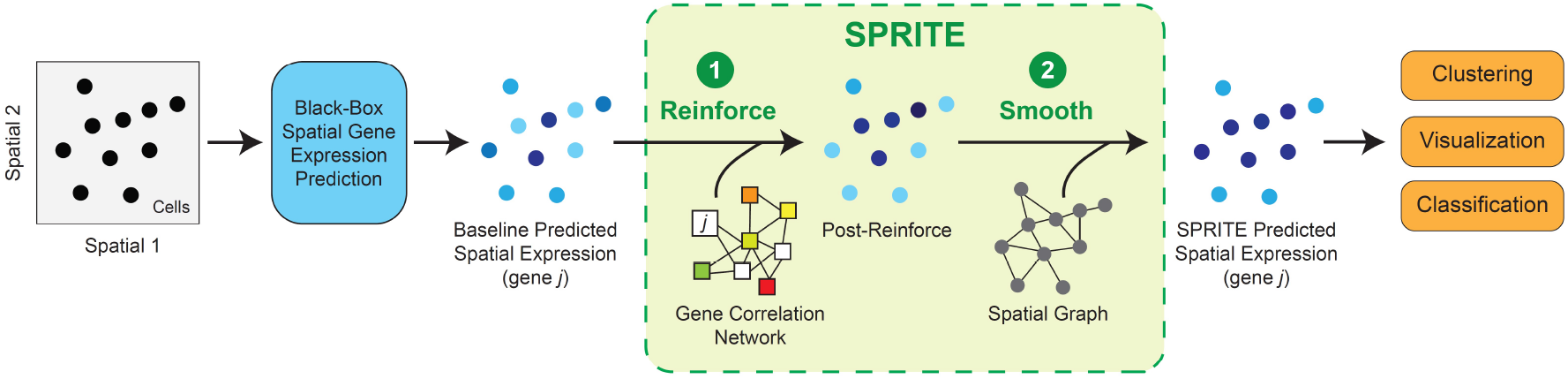
Illustration of the steps in the SPRITE algorithm. Baseline spatial gene expression predictions are made from any available prediction methods. SPRITE builds a gene correlation network for the Reinforce step to propagate prediction errors from calibration genes to unseen genes. SPRITE builds a spatial graph of neighboring cells for the Smooth step to propagate predicted expression across neighbors. The SPRITEprocessed predicted expression can be used in downstream analysis tasks.

#### 2.3.1 Reinforce step with gene correlation network

In the Reinforce step, we use a modified version of the iterative smoothing procedure [28, 29]. Specifically, the update rule for Reinforce is:

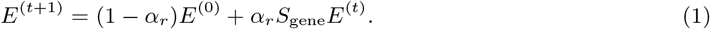

In Eq. (1), *E*^(0)^ = *E* is the initial residuals matrix containing the prediction errors, *E*^(*t*)^ is the residuals matrix after the *t*-th update, *S*_gene_ is a normalized adjacency matrix from a gene correlation network, and *α*_*r*_ is the smoothing parameter for Reinforce. The residuals matrix is computed as *E* = (*Y* − *G*)^*T*^, where *G* is the predicted gene expression matrix and *Y* is the measured gene expression in *X*_spatial_ masked for calibration genes. We set *E* = 0 for all target genes. To select a value for *α*_*r*_, we used nested cross-validation where we held out different subsets of calibration genes by setting *E* = 0 for those genes, performed Reinforce until convergence, and then computed the mean absolute error (MAE) of the propagated residuals with respect to the ground truth residuals of the held-out calibration genes. We selected the *α*_*r*_ corresponding to the minimum average MAE across all calibration genes. We performed this search over 10 uniformly spaced values of *α*_*s*_ ranging from 0.01 to 0.9. Using the predicted spatial gene expression as input, we iterate the Reinforce update rule until convergence when *E*^(*t*+1)^ = *E*^(*t*)^. Across all benchmark spatial transcriptomics datasets, the Reinforce step generally converged in less than 100 iterations.

The Reinforce step propagates residuals along a gene correlation network, defined by the normalized adjacency matrix *S*_gene_. We build the gene correlation network according to the following procedure: (1) calculate pairwise Spearman rank correlation between predicted expression of all genes in the combined target and calibration gene sets (*r* genes in total); (2) automatically compute a threshold Spearman rank correlation for drawing edges between genes such that all genes have at least one neighbor (i.e. the graph is connected); (3) apply the threshold to build an adjacency matrix with binary values indicating existence of edges between nodes. The adjacency matrix is then normalized by dividing the elements of each row by the row sum to create *S*_gene_ ∈ ℝ^*r×r*^. In this procedure, the thresholds are specific to each spatial transcriptomics dataset and selected to ensure that information can be propagated across the full network. Across all benchmark spatial transcriptomics datasets, the automatic thresholds for the Spearman rank correlation generally ranged from 0.05 to 0.4 for Harmony-kNN, from 0.15 to 0.55 for SpaGE, and from 0.5 to 0.9 for Tangram.

In the final step of the Reinforce step, after convergence of Eq. (1) has been reached, we compute the reinforced spatial gene expression prediction by adding the reinforced residual matrix *E*^(*t*)^ to the initial predictions: *G*^(0)^ = *G* + *E*^(*t*)^.

#### 2.3.2 Smooth step with spatial neighborhood graph

In the Smooth step, we use a similar iterative smoothing procedure as in the Reinforce step. Specifically, the update rule for Smooth is:

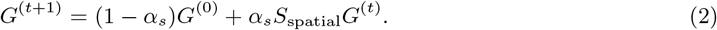

In Eq. (2), *G*^(0)^ is the reinforced spatial gene expression prediction matrix, *G*^(*t*)^ is the spatial gene expression matrix after the *t*-th update, *S*_spatial_ is a normalized adjacency matrix from a spatial neighborhood graph of the cells in the spatial transcriptomics data, and *α*_*s*_ is the smoothing parameter for Smooth. The input to the Smooth step is the output of the Reinforce step, *G*^(0)^ = *G* + *E*^(*t*)^. For the ablated SPRITE model without the Reinforce step (No Reinforce), we set *G*^(0)^ = *G*. To select the optimal *α*_*s*_, we performed a line search across 10 uniformly spaced values of *α*_*s*_ ranging from 0.01 to 0.9, computing the MAE between the smoothed predictions outputted by the Smooth step and the measured expression for all calibration genes. We selected the value of *α*_*s*_ that minimized this MAE measure. Using the reinforced spatial gene expression predictions as input, we iterate the Smooth update rule until convergence when *G*^(*t*+1)^ = *G*^(*t*)^. Across all benchmark spatial transcriptomics datasets, the Smooth step generally converged in less than 100 iterations.

To construct spatial neighborhood graphs with nodes corresponding to the cells in *X*_spatial_, we drew edges between each cell and its *k* nearest neighbors according to Euclidean distance between the centroids of each cell. We set *k* = 50 for all analyses presented in this study, but also observed that the performance of SPRITE was generally robust to most reasonable choices of *k*. To remove outlier edges that span exceptionally large distances, we removed all edges with distance greater than 1.5 times the interquartile range of all edge distances. This resulted in a binary adjacency matrix representing the edges in the spatial neighborhood graph. Importantly, propagation of predicted gene expression across cells of different cell types or subtypes is not desirable. To selectively propagate information to cells with similar transcriptomic profiles (i.e. similar cell type), we weighted each edge by the similarity in measured ground truth expression of all genes in the spatial transcriptomics data between neighboring cells. To reduce distortions from high-dimensional vectors, we first applied principal component analysis to the measured gene expression and used the top five principal components to compute the cosine similarity for weighting each edge. We then normalized the weighted adjacency matrix by dividing each row by its sum to generate *S*_spatial_. Finally, the predicted gene expression values are propagated across the spatial neighborhood graph according to Eq. (2) until convergence. The output of the Smooth step *G*^(*t*)^ is the SPRITE predicted spatial gene expression matrix.

### 2.4 Experiments and evaluation

#### 2.4.1 Improvement in prediction

To evaluate SPRITE across the benchmark datasets and spatial gene expression prediction methods, we employed several statistical metrics to reflect performance in prediction quality. For a given gene, we measured the Pearson correlation coefficient (PCC) and the mean absolute error (MAE) between its baseline predicted expression and its measured expression across all cells in the spatial transcriptomics data. In the same gene, we also measured the PCC and MAE for the SPRITE predicted expression and the measured expression. To compute the improvement in performance provided by SPRITE, we computed the difference of each measure between the baseline predicted expression and the SPRITE predicted expression. The difference was computed such that positive values indicated either higher PCC or lower MAE for the SPRITE predictions.

#### 2.4.2 Cell clustering and evaluation

We performed clustering of cells using the Leiden algorithm [30]. First, we set all negative values to zero in the gene expression matrix, normalized and log-transformed the counts, and performed principal component analysis (PCA). Then, we performed Leiden clustering using the default settings in the Scanpy package [31] (version 1.9.3).

For the clustering experiments, we used the three spatial transcriptomics datasets that had publicly available cell type annotations, which included the mouse somatosensory osmFISH dataset, the mouse gastrulation seqFISH dataset, and the axolotl telencephalon Stereo-seq dataset. For the mouse somatosensory osmFISH dataset, we used cell type annotations under ‘ClusterName’ from the metadata available at http://linnarssonlab.org/osmFISH/osmFISH_SScortex_mouse_all_cells.loom. For the mouse gastrulation seqFISH dataset, we used cell type annotations under ‘cell type mapped refined’ from the metadata available at https://content.cruk.cam.ac.uk/jmlab/SpatialMouseAtlas2020/ in ‘metadata.Rds’ file for ‘embryo1’ and ‘z5’. For the axolotl telencephalon Stereo-seq dataset, we retrieved cell type annotations under ‘Annotation’ from the metadata available at https://db.cngb.org/stomics/artista/ for the ‘Stage44.h5ad’ object file.

To evaluate the quality of cell clusters, we used the adjusted Rand index to compare cluster assignments obtained on either baseline predicted spatial gene expression or SPRITE predicted spatial gene expression to the ground truth cell type labels.

#### 2.4.3 Cell visualization and evaluation

We generated two-dimensional visualizations of cells in spatial transcriptomics data using the UMAP [32], t-SNE [33] with PCA initialization, and PCA algorithms applied to different versions of the gene expression matrix. The gene expression matrices were standardized by features before input to the visualization algorithms. We performed visualization with all three algorithms on all eleven benchmark datasets.

To evaluate the quality of these visualizations, we computed the Spearman visualization score, which is the Spearman rank correlation coefficient between the pairwise Euclidean distances of cells in the visualization coordinates and in the original high-dimensional data. The metric was computed using the concordance score method in DynamicViz [34].

#### 2.4.4 Cell type classification and evaluation

We used L1-penalized logistic regression models and trained them on different versions of the gene expression matrix to predict cell type labels (using up to the three most prevalent cell types). The models were evaluated under stratified 5-fold cross-validation using the cell type labels for stratification. To evaluate the quality of the classifiers, we computed the accuracy, macro-averaged F1 score, and the area under the receiver operating characteristic curve (AUC-ROC). These metrics were averaged across all cross-validation folds.

## 3 Results

### 3.1 SPRITE improves prediction of spatial gene expression

We developed SPRITE, a meta-algorithm that post-processes predicted spatial gene expression by propagating prediction errors across a gene correlation network (Reinforce) and smooths predictions across a spatial neighborhood graph (Smooth) (Fig. 1). To evaluate whether SPRITE improved the quality of spatial gene expression predictions, we applied SPRITE to spatial gene expression predictions generated from three methods (SpaGE, Tangram, Harmony-kNN) and across eleven paired spatial transcriptomics and RNAseq dataset benchmarks. Relative to the baseline predicted spatial gene expression, SPRITE predicted spatial gene expression was generally better correlated and had lower mean absolute error with respect to the measured ground truth expression (Fig. 2AB). Interestingly, for Tangram, SPRITE produced slightly negative improvement under the correlation metric but large positive improvement under the error metric (Fig. 2AB). This trend was due to the tendency of Tangram to predict extreme values for gene expression, which can skew correlation measures but also provides opportunity for large reductions in prediction error. In aggregate, all evaluation contexts (i.e. unique combination of dataset and prediction method) observed improvements in at least one of the aforementioned performance metrics for prediction quality (Fig. 2C). Moreover, most genes also observed improvements in at least one of the performance metrics (Fig. 2C). In many cases, the improvement in gene expression prediction when using SPRITE is visually striking. For example, SPRITE can recover spatial expression patterns that were missing in the baseline predictions (Fig. 2D). Conversely, SPRITE can also attenuate high baseline predictions to better capture selective spatial expression of certain genes (Fig. 2E). In summary, SPRITE provides general improvements in the quality of predicted spatial gene expression across different datasets and prediction methods.

**Figure 2:**
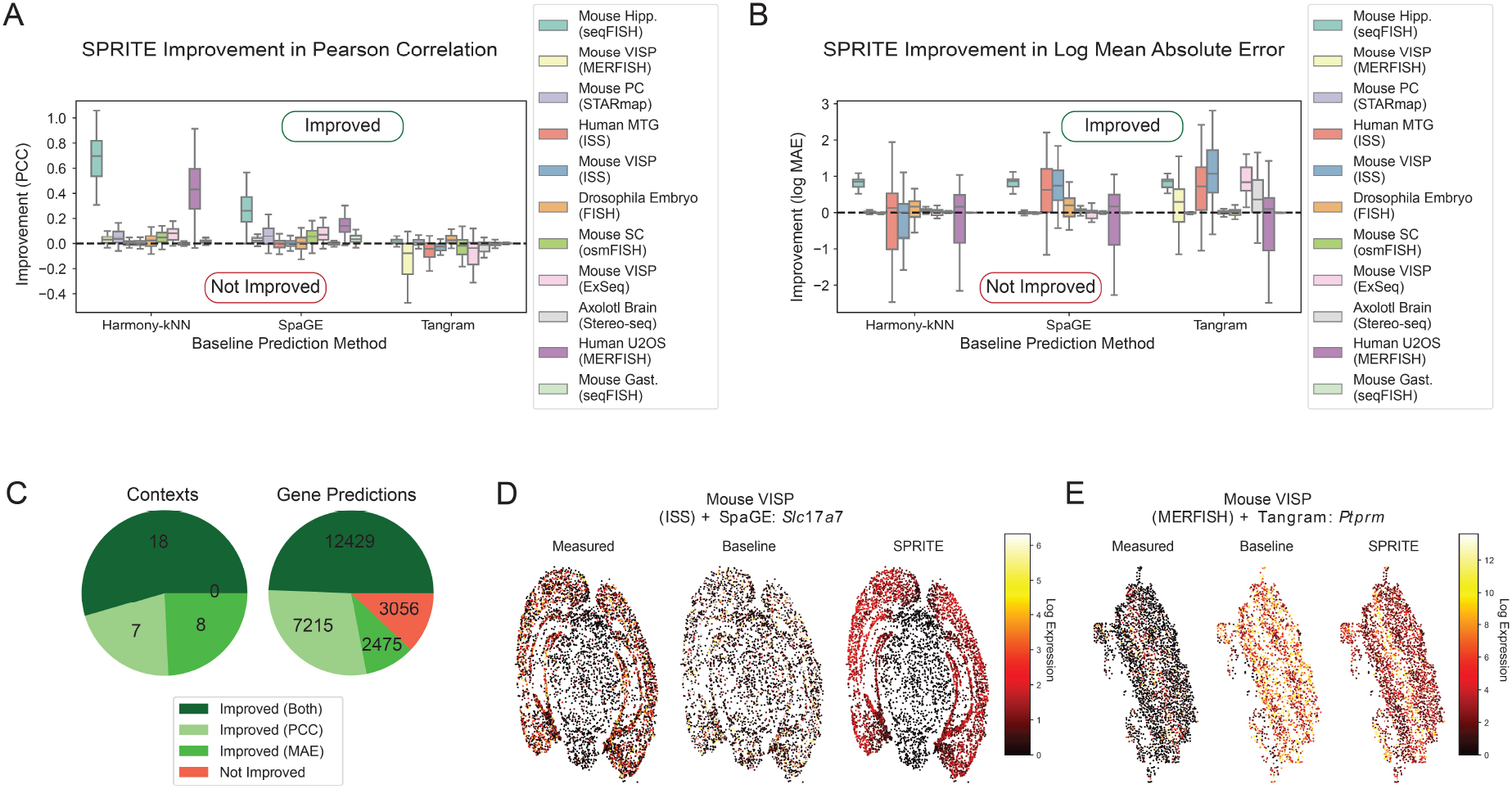
SPRITE results in improved spatial gene expression prediction. (A-B) SPRITE improvements over baseline spatial gene expression prediction across eleven spatial transcriptomics and single-cell RNAseq dataset pairs and three spatial baseline prediction methods (SpaGE, Tangram, Harmony-kNN) as measured by gene-wise prediction quality metrics: (A) the Pearson correlation coefficient and (B) the log mean absolute error between the predicted and measured spatial gene expression. Improvement is calculated as the difference in the metrics for the baseline predicted expression and the corresponding SPRITE predicted expression. (C) Pie graphs illustrating the breakdown of improvements in predicted spatial gene expression by context (dataset and prediction method pair) (left) and by individual genes (right). (D-E) Spatial visualization of cells colored by (from left to right): the normalized measured gene expression, baseline predicted gene expression, and SPRITE predicted gene expression for (D) the transcript *Slc17a7* in the ISS dataset of mouse visual cortex using SpaGE for baseline prediction and (E) the transcript *Ptprm* in the MERFISH dataset of mouse visual cortex using Tangram for baseline prediction.

### 3.2 Reinforce and Smooth steps are necessary for improvements by SPRITE

Given the improvement in predicted gene expression provided by SPRITE, we investigated the relative contributions of the Reinforce and Smooth steps to the observed improvement. To do so, we developed SPRITE implementations where one of the two steps was ablated from the pipeline (i.e. “No Reinforce” and “No Smooth”). After ablating these steps, we performed identical experiments as before to measure the improvement in prediction provided through the modified versions of SPRITE. We observed that both Smooth and Reinforce can provide improvements over baseline prediction when used in isolation, but that additional improvements can be achieved when they are combined such as in SPRITE (Fig. 3AB). This result suggests that Reinforce and Smooth are synergistic in the SPRITE algorithm.

**Figure 3:**
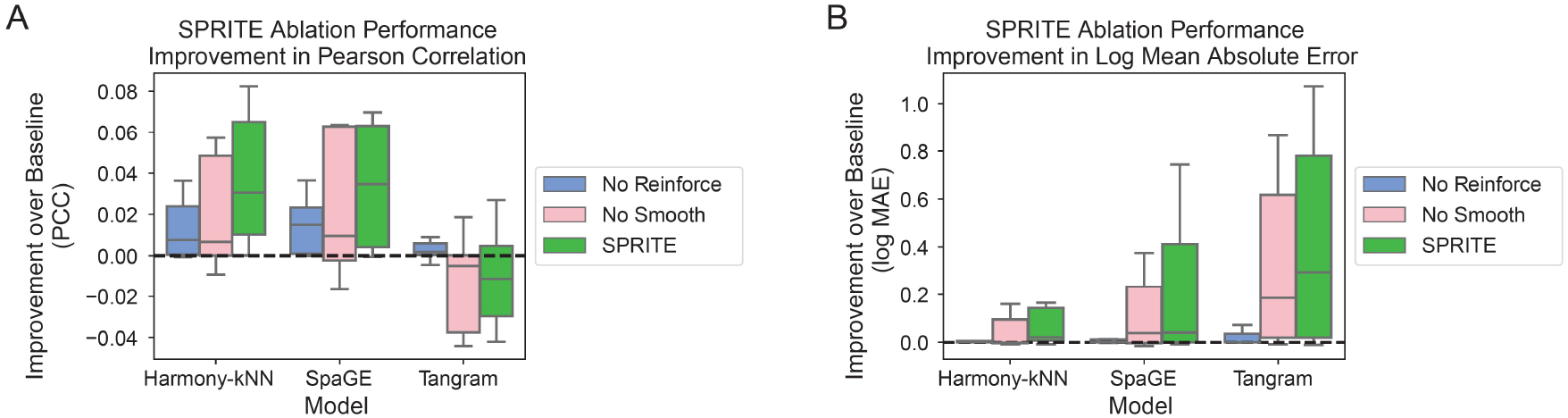
Ablation of Reinforce or Smooth steps reduce SPRITE performance. (A-B) Improvement over baseline spatial gene expression prediction for different post-processing methods (SPRITE, ablation of Reinforce step, ablation of Smooth step) as measured by gene-wise prediction quality metrics: (A) the Pearson correlation coefficient and (B) the log mean absolute error between the predicted and measured spatial gene expression. Results are aggregated across eleven spatial transcriptomics and RNAseq dataset pairs and three spatial baseline prediction methods (SpaGE, Tangram, Harmony-kNN).

### 3.3 SPRITE improves clustering of cells by cell type

Of particular interest is whether the improvements in prediction quality by SPRITE can be transferred to common downstream analysis tasks such as the clustering of cells. Clustering is often used to identify cell types or subtypes in single-cell transcriptomics. We compared Leiden clustering assignments of the baseline predicted spatial gene expression to those of the SPRITE predicted spatial gene expression (Fig. 4A). Across three spatial transcriptomics datasets with cell type annotations and three prediction methods, the cluster assignments obtained from the SPRITE predicted spatial gene expression produced strictly better concordance with the ground truth labels than the cluster assignments from the baseline predicted gene expression, as measured by the adjusted Rand index (Fig. 4B). SPRITE improves clustering of cells by predicted gene expression to better capture underlying cell type information.

**Figure 4:**
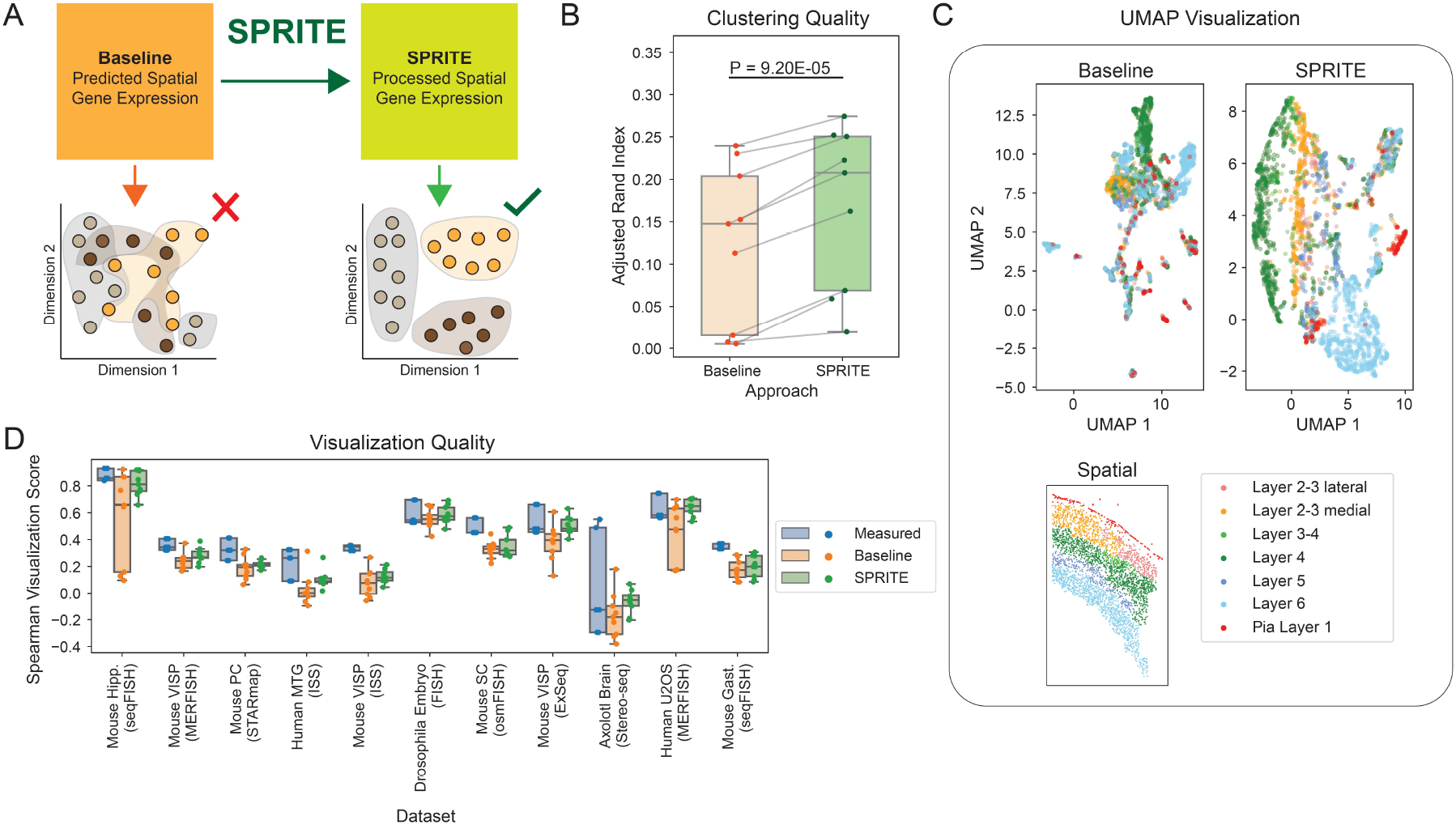
SPRITE predicted spatial gene expression provides more faithful clustering and visualization of cells. (A) Illustration of evaluation scheme where cells are represented by clustering and visualization based on either baseline predicted spatial transcriptomes or SPRITE-predicted transcriptomes. (B) Cell clustering quality measured by the adjusted Rand index with cell type as the ground truth label on all three spatial transcriptomics datasets that included cell type annotations. Clustering used the Leiden algorithm on either baseline predicted spatial gene expression or SPRITE predicted spatial gene expression. (C) Visualization of all cells in the osmFISH mouse somatosensory cortex dataset using UMAP projections of either the baseline predicted spatial gene expression (top left) or the SPRITE predicted spatial gene expression (top right) along with a cutout showing the spatial location of all cells (bottom). Cells are colored by their anatomic region labels. (D) Spearman visualization scores indicating preservation of pairwise cell distances in two-dimensional visualization with respect to the original high-dimensional data using either the measured gene expression, baseline predicted gene expression, or SPRITE predicted gene expression profiles for visualization. Results are shown for eleven spatial transcriptomics datasets and aggregated over three spatial gene expression prediction methods (SpaGE, Tangram, Harmony-kNN) and three dimensionality reduction methods for visualization (UMAP, t-SNE, PCA).

### 3.4 SPRITE improves visualization of predicted spatial transcriptomics

We further investigated the performance of SPRITE in data visualization, which is often used to intuitively understand high-dimensional data such as spatial transcriptomics. We used dimensionality reduction algorithms to obtain low-dimensional visualizations of the measured ground truth, baseline predicted, and SPRITE predicted spatial gene expression. We visualized the mouse somatosensory osm-FISH dataset using UMAP on the baseline and SPRITE predicted expression, and the visualizations with SPRITE showed markedly better separation of cells by their anatomic region, which is desirable since the tissue is highly structured (Fig. 4C). More broadly, we observed that visualizations (UMAP, t-SNE, and PCA) generated on SPRITE predicted spatial gene expression were consistently better than visualizations obtained from baseline predicted spatial gene expression at preserving the pairwise distances between cells with respect to the original high-dimensional data and were often comparable in quality to visualizations generated from the measured ground truth gene expression (Fig. 4D). As such, visualizations of SPRITE predicted expression can more faithfully represent the spatial location of cells and their relations in the spatial transcriptomics data.

### 3.5 SPRITE improves cell type classification

Finally, we explored whether SPRITE would improve underlying biological signal in the predicted expression that could be detected by models trained and evaluated on the data. Specifically, we compared the performance of cell type classifiers trained on either baseline predicted expression or on SPRITE predicted expression (Fig. 5A). Across the three spatial transcriptomics datasets with cell type annotations and evaluated on three different classification metrics, we observed consistent and statistically significant improvements in classification performance when models are trained on the SPRITE predicted expression (Fig. 5B-D). In several cases, the performance of the SPRITE-based model can even exceed that of models trained on the measured expression (Fig. 5B-D).

**Figure 5:**
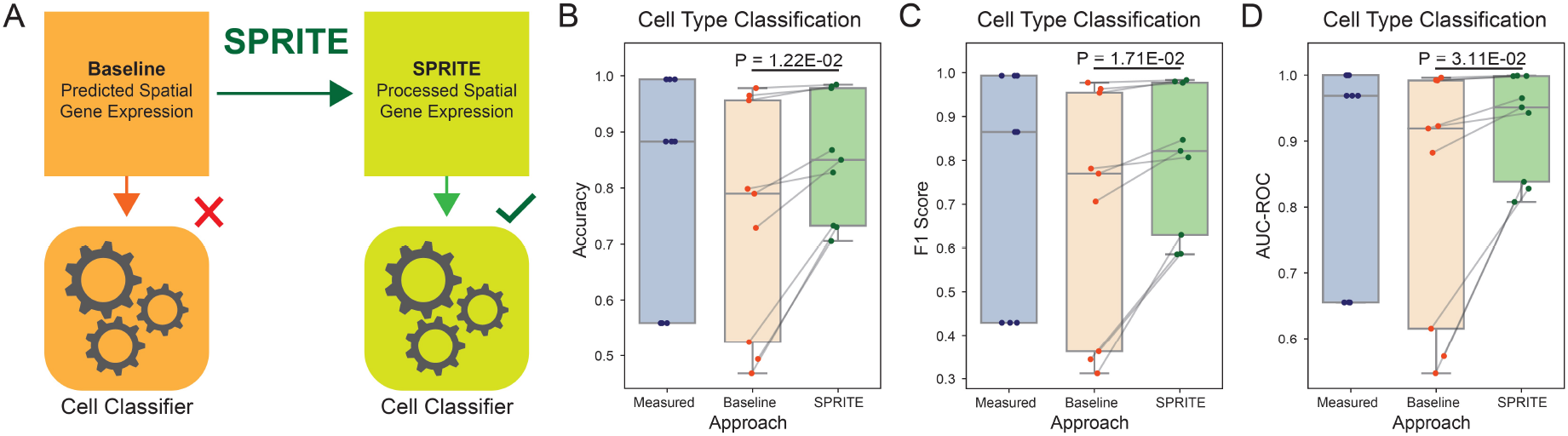
SPRITE predicted spatial gene expression provides better signal for cell type classification. (A) Illustration of the evaluation scheme for training classifiers to predict cell type using baseline predicted spatial transcriptomes and SPRITE-predicted transcriptomes. (B-D) Cell type classification performance metrics for classifiers trained and evaluated (5-fold cross-validation) on the measured spatial gene expression, baseline predicted spatial gene expression, and the SPRITE predicted spatial gene expression: (B) accuracy of cell type classification, (C) F1 score of cell type classification, and (D) area under the receiver-operator curve (AUC-ROC) of cell type classification. Results were aggregated across three spatial transcriptomics datasets with cell type labels and three spatial gene expression prediction methods (SpaGE, Tangram, Harmony-kNN). P-values were computed using a paired two-sided t-test.

## 4 Discussion

SPRITE is a meta-algorithm that functions with any existing or future spatial gene expression prediction method. We show that SPRITE generally increases the quality of spatial gene expression predictions, and that this improvement is the combined result of the Reinforce and Smooth steps in the SPRITE pipeline. Furthermore, we show that SPRITE can be extended to improve common downstream analysis tasks such as clustering of cells, faithful visualization of spatial transcriptomics, and classification of cells by cell type. Surprisingly, in several tasks, SPRITE not only outperforms baseline methods but can exceed approaches that use ground truth measured expression, suggesting that in addition to improved spatial gene expression prediction, SPRITE may also confer other advantages perhaps through de-noising gene expression. Ultimately, we believe that SPRITE will be a valuable tool in leveraging spatial transcriptomics to uncover new biological processes.

SPRITE is highly scalable since its computational complexity is related by a multiplicative constant to that of the underlying prediction method responsible for generating the baseline predicted gene expression. This constant is determined by the number of cross-validation folds for calibration gene expression prediction and can therefore be adjusted for different contexts and desired runtimes. The SPRITE runtime is generally dominated by the generation of baseline spatial gene expression prediction. For example, the median single-threaded runtimes for SPRITE across the benchmark datasets were 242 seconds for generating 10-fold spatial gene expression predictions, 13 seconds for Reinforce, and 33 seconds for Smooth. Since our ablation studies showed that both Reinforce and Smooth can independently provide improvements over baseline predictions, in cases demanding particularly high computational efficiency, SPRITE (No Reinforce) may be a desirable alternative to the full SPRITE algorithm since the Smooth step does not require generating multiple cross-validation folds for predicting calibration gene expression. An open question is whether SPRITE can still improve predicted gene expression if the underlying baseline prediction method explicitly uses spatial information and gene correlation network information. Currently, no baseline spatial gene expression prediction method meets both criteria. Similarly, it would be worthwhile to design end-to-end prediction methods that use spatial and gene correlation information and compare the performance of these methods to the multi-step approach used by SPRITE.

Although we have considered several common spatial transcriptomics analysis tasks, exciting new approaches are emerging in this field for investigating cellular interactions and characterizing local neighborhood effects [1]. Testing the ability of SPRITE to yield improvements on these additional frameworks would further extend the range of applications for SPRITE and spatial gene expression prediction. Additionally, combining SPRITE with new methods for estimating spatial gene expression prediction uncertainty [26] may provide an ecosystem for interpreting conclusions drawn from predicted spatial gene expression and performing useful scientific inference.

## Author contributions statement

E.D.S. conceived of the method with input from J.Z. and R.M., E.D.S. conducted the experiments and analyzed the results, E.D.S. wrote the manuscript with input from J.Z. and R.M.

## Acknowledgments

Funding support was provided by Knight-Hennessy Scholars program (E.D.S.), Paul and Daisy Soros Fellowship for New Americans (E.D.S.), the National Science Foundation Graduate Research Fellowship Program (E.D.S.), Professor David Donoho at Stanford University (R.M.), NSF CAREER 1942926 (J.Z.), NIH P30AG059307 (J.Z.), 5RM1HG010023 (J.Z.) and grants from the Silicon Valley Foundation (J.Z.) and the Chan-Zuckerberg Initiative (J.Z.).

## Code availability

The SPRITE software package is available at https://github.com/sunericd/SPRITE. Code for generating experiments and analyses in the manuscript is available at https://github.com/sunericd/sprite-figures-and-analyses.

## References

[1] Moses, L. & Pachter, L. Museum of spatial transcriptomics. Nature Methods 1–13 (2022). URL https://www.nature.com/articles/s41592-022-01409-2. Publisher: Nature Publishing Group.

[2] Li, B. et al. Benchmarking spatial and single-cell transcriptomics integration methods for transcript distribution prediction and cell type deconvolution. Nature Methods 19, 662–670 (2022). URL https://www.nature.com/articles/s41592-022-01480-9. Number: 6 Publisher: Nature Publishing Group.

[3] Abdelaal, T., Mourragui, S., Mahfouz, A. & Reinders, M. J. T. SpaGE: Spatial Gene Enhancement using scRNA-seq. Nucleic Acids Research 48, e107 (2020). URL 10.1093/nar/gkaa740.

[4] Allen, W. E., Blosser, T. R., Sullivan, Z. A., Dulac, C. & Zhuang, X. Molecular and spatial signatures of mouse brain aging at single-cell resolution. Cell 186, 194–208.e18 (2023). URL https://www.cell.com/cell/abstract/S0092-8674(22)01523-9. Publisher: Elsevier.

[5] Shengquan, C., Boheng, Z., Xiaoyang, C., Xuegong, Z. & Rui, J. stPlus: a reference-based method for the accurate enhancement of spatial transcriptomics. Bioinformatics 37, i299–i307 (2021). URL 10.1093/bioinformatics/btab298.

[6] Welch, J. D. et al. Single-Cell Multi-omic Integration Compares and Contrasts Features of Brain Cell Identity. Cell 177, 1873–1887.e17 (2019).

[7] Biancalani, T. et al. Deep learning and alignment of spatially resolved single-cell transcriptomes with Tangram. Nature Methods 18, 1352–1362 (2021). URL https://www.nature.com/articles/s41592-021-01264-7. Number: 11 Publisher: Nature Publishing Group.

[8] Long, B., Miller, J. & Consortium, T. S. SpaceTx: A Roadmap for Benchmarking Spatial Transcriptomics Exploration of the Brain (2023). URL http://arxiv.org/abs/2301.08436. ArXiv:2301.08436 [q-bio].

[9] Joglekar, A. et al. A spatially resolved brain region- and cell type-specific isoform atlas of the postnatal mouse brain. Nature Communications 12, 463 (2021).

[10] Booeshaghi, A. S. et al. Isoform cell-type specificity in the mouse primary motor cortex. Nature 598, 195–199 (2021). URL https://www.nature.com/articles/s41586-021-03969-3. Number: 7879 Publisher: Nature Publishing Group.

[11] Wang, X. et al. Three-dimensional intact-tissue sequencing of single-cell transcriptional states. Science 361, eaat5691 (2018). URL 10.1126/science.aat5691. Publisher: American Association for the Advancement of Science.

[12] Gyllborg, D. et al. Hybridization-based in situ sequencing (HybISS) for spatially resolved transcriptomics in human and mouse brain tissue. Nucleic Acids Research 48, e112 (2020). URL 10.1093/nar/gkaa792.

[13] Karaiskos, N. et al. The Drosophila embryo at single-cell transcriptome resolution. Science 358, 194–199 (2017). URL 10.1126/science.aan3235. Publisher: American Association for the Advancement of Science.

[14] Nitzan, M., Karaiskos, N., Friedman, N. & Rajewsky, N. Gene expression cartography. Nature 576, 132–137 (2019). URL https://www.nature.com/articles/s41586-019-1773-3. Number: 7785 Publisher: Nature Publishing Group.

[15] Codeluppi, S. et al. Spatial organization of the somatosensory cortex revealed by osmFISH. Nature Methods 15, 932–935 (2018). URL https://www.nature.com/articles/s41592-018-0175-z. Number: 11 Publisher: Nature Publishing Group.

[16] Alon, S. et al. Expansion sequencing: Spatially precise in situ transcriptomics in intact biological systems. Science 371, eaax2656 (2021). URL 10.1126/science.aax2656. Publisher: American Association for the Advancement of Science.

[17] Hodge, R. D. et al. Conserved cell types with divergent features in human versus mouse cortex. Nature 573, 61–68 (2019). URL https://www.nature.com/articles/s41586-019-1506-7. Number: 7772 Publisher: Nature Publishing Group.

[18] Yao, Z. et al. A taxonomy of transcriptomic cell types across the isocortex and hippocampal formation. Cell 184, 3222–3241.e26 (2021). URL https://www.sciencedirect.com/science/article/pii/S0092867421005018.

[19] Tasic, B. et al. Shared and distinct transcriptomic cell types across neocortical areas. Nature 563, 72–78 (2018). URL https://www.nature.com/articles/s41586-018-0654-5. Number: 7729 Publisher: Nature Publishing Group.

[20] Shah, S., Lubeck, E., Zhou, W. & Cai, L. In Situ Transcription Profiling of Single Cells Reveals Spatial Organization of Cells in the Mouse Hippocampus. Neuron 92, 342–357 (2016). URL https://www.sciencedirect.com/science/article/pii/S0896627316307024.

[21] Lohoff, T. et al. Integration of spatial and single-cell transcriptomic data elucidates mouse organogenesis. Nature Biotechnology 40, 74–85 (2022). URL https://www.nature.com/articles/s41587-021-01006-2. Number: 1 Publisher: Nature Publishing Group.

[22] Wei, X. et al. Single-cell Stereo-seq reveals induced progenitor cells involved in axolotl brain regeneration. Science 377, eabp9444 (2022). URL 10.1126/science.abp9444. Publisher: American Association for the Advancement of Science.

[23] Lust, K. et al. Single-cell analyses of axolotl telencephalon organization, neurogenesis, and regeneration. Science 377, eabp9262 (2022). URL 10.1126/science.abp9262. Publisher: American Association for the Advancement of Science.

[24] Zhou, Y. et al. Single-cell RNA landscape of intratumoral heterogeneity and immunosuppressive microenvironment in advanced osteosarcoma. Nature Communications 11, 6322 (2020).

[25] Xia, C., Fan, J., Emanuel, G., Hao, J. & Zhuang, X. Spatial transcriptome profiling by MERFISH reveals subcellular RNA compartmentalization and cell cycle-dependent gene expression. Proceedings of the National Academy of Sciences 116, 19490–19499 (2019). URL 10.1073/pnas.1912459116. Publisher: Proceedings of the National Academy of Sciences.

[26] Sun, E. D., Ma, R., Navarro Negredo, P., Brunet, A. & Zou, J. TISSUE: uncertainty-calibrated prediction of single-cell spatial transcriptomics improves downstream analyses. bioRxiv 2023.04.25.538326 (2023). URL https://www.ncbi.nlm.nih.gov/pmc/articles/PMC10168375/.

[27] Korsunsky, I. et al. Fast, sensitive and accurate integration of single-cell data with Harmony. Nature Methods 16, 1289–1296 (2019). URL https://www.nature.com/articles/s41592-019-0619-0. Number: 12 Publisher: Nature Publishing Group.

[28] Zhou, D., Bousquet, O., Lal, T., Weston, J. & Schölkopf, B. Learning with Local and Global Consistency. In Thrun, S., Saul, L. & Schölkopf, B. (eds.) Advances in Neural Information Processing Systems, vol. 16 (MIT Press, 2003). URL https://proceedings.neurips.cc/paper/2003/file/87682805257e619d49b8e0dfdc14affa-Paper.pdf.

[29] Huang, Q., He, H., Singh, A., Lim, SN. & Benson, A. R. Combining Label Propagation and Simple Models Out-performs Graph Neural Networks (2020). URL http://arxiv.org/abs/2010.13993. ArXiv:2010.13993 [cs].

[30] Traag, V. A., Waltman, L. & van Eck, N. J. From Louvain to Leiden: guaranteeing well-connected communities. Scientific Reports 9, 5233 (2019). URL https://www.nature.com/articles/s41598-019-41695-z. Number: 1 Publisher: Nature Publishing Group.

[31] Wolf, F. A., Angerer, P. & Theis, F. J. SCANPY: large-scale single-cell gene expression data analysis. Genome Biology 19, 15 (2018). URL 10.1186/s13059-017-1382-0.

[32] McInnes, L., Healy, J., Saul, N. & Großberger, L. UMAP: Uniform Manifold Approximation and Projection. Journal of Open Source Software 3, 861 (2018). URL 10.21105/joss.00861.

[33] van der Maaten, L. J. P. & Hinton, G. E. Visualizing High-Dimensional Data Using t-SNE. Journal of Machine Learning Research 9, 2579–2605 (2008). Publisher: Microtome Publishing.

[34] Sun, E. D., Ma, R. & Zou, J. Dynamic visualization of high-dimensional data. Nature Computational Science 3, 86–100 (2023). URL https://www.nature.com/articles/s43588-022-00380-4. Number: 1 Publisher: Nature Publishing Group.

